# Visual Representation of Experimental Protocols

**DOI:** 10.1101/226852

**Authors:** James Scott-Brown, Antonis Papachristodoulou

## Abstract

Using robots to automate laboratory tasks could increase throughput and reproducibility, but requires experimental protocols to be specified in a computer-readable format. We present a new user interface (Lists of Liquids) for specifying experimental protocols by directly manipulating a diagram: rather than having to specify individual liquid handling operations, the user can simply specify that particular lists of liquids should be combined as either a Cartesian product or convolution, and the system will plan a series of liquid handling steps to achieve this. This is intended to provide a higher-level interface in order to make the creation of protocols faster and less error prone.

## 1 Introduction

There have been various attempts at constructing formal languages for expressing experimental protocols. These are intended to reduce ambiguity compared to natural language and enable either a robot to automatically execute a protocol, or a computer to guide a human operator through a protocol. These include both textual interfaces (e.g. BioCoder (1), PaR-PaR (2), Aquarium (3), Autoprotocol (4), Antha (5), and Symbolic Lab Language (6)) and graphical interfaces (e.g. BioBlocks (7), Wet Lab Accelerator (8), and proprietary tools from robotic systems manufacturers).

However, these interfaces place the primary focus on the specific liquid handling steps to be performed, rather than their intended results. We present here a graphical interface that reverses this focus: the user expresses what they want to achieve at a high level, and the system determines what movement of liquids between physical locations are necessary to achieve this. The system can produce a range of outputs, including a diagram of the specified operations, a description of the protocol in English, and an executable protocol in various formats (including Autoprotocol-python and OpenTrons).

In contrast to previous tools, this is a single open-source application that provides one interface that can be used to control a range of automation platforms.

The decision to make previous interfaces graphical was generally motivated by a desire to avoid requiring users to be familiar with any programming language (7, 8). However, the graphical representation that we have chosen has the additional advantage of producing a diagram with a structure reflecting the protocol’s structure, rather than just representing it as a linear sequence of steps; this makes it easier to understand what a protocol is intended to do or identify errors.

**Figure 1:**
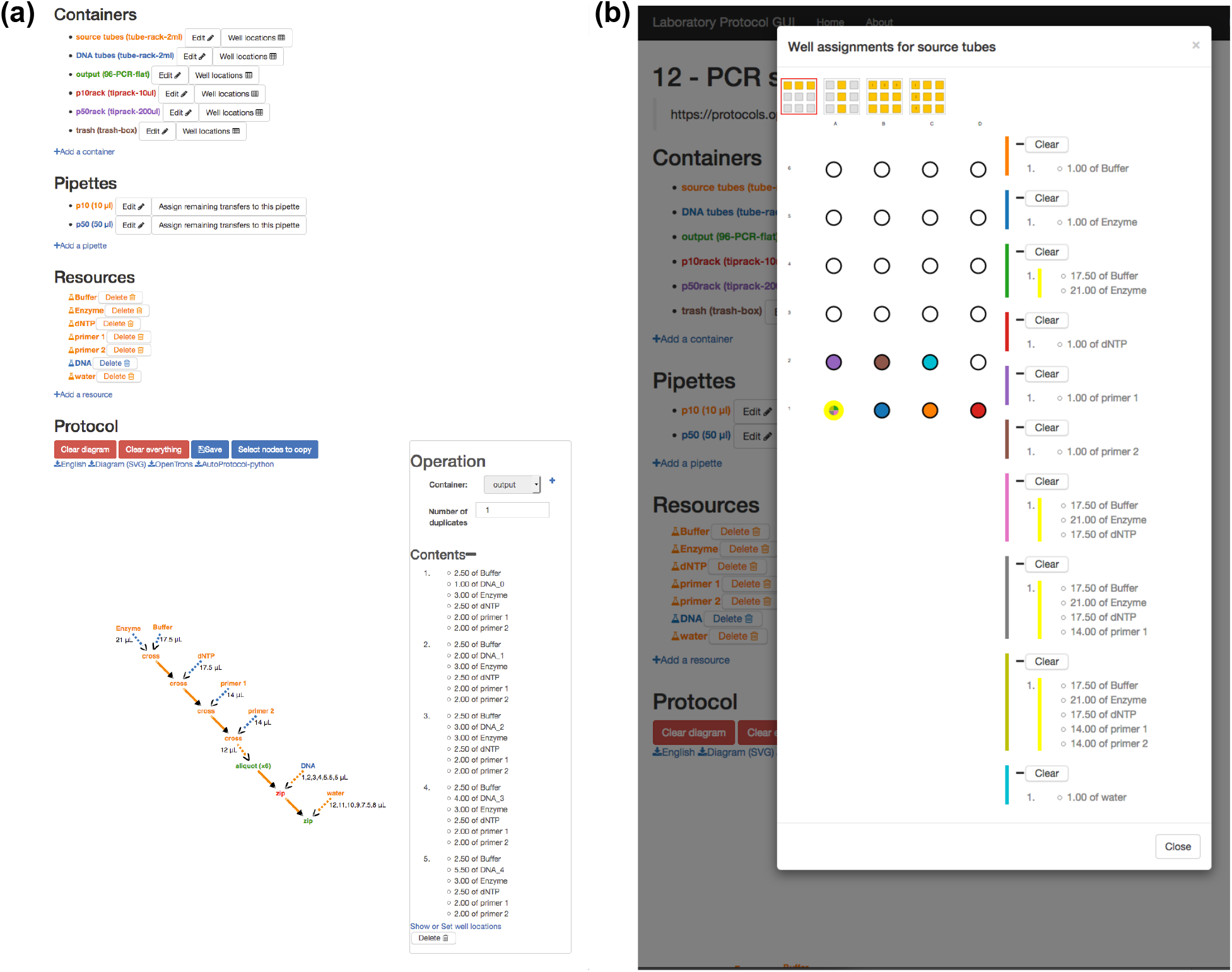
Screenshot showing the List of Liquids interface. **Left**: a list of the containers, pipettes, and resources required by the protocol, followed by a diagram showing the protocol’s operations steps. A node labeled zip is highlighted in red, and a panel to the right shows its details, including a list of its contents. **Right**: modal dialog for assigning liquids to specific wells on a container. The cursor is hovering over one well, causing both it and the corresponding entries in the list on the right (which list its contents at different stages of the protocol) to be highlighted in yellow.

## 2 Diagrammatic representation of protocols

A protocol is represented by a list of required things (containers, pipettes, resources), and a diagram showing how they are used.

- *Containers* are things that can contain liquids (or tip-racks, which contain disposable tips for pipettes, rather than liquids). Some container types consist of a single compartment (e.g. a trash container), whereas others consist of multiple wells (e.g. a 96-well plate).
- *Pipettes* are the pipettes used to physically transfer liquids between locations. To export a protocol in OpenTrons format, the pipette used for each transfer must be specified, but this is not necessary for all output formats.
- *Resources* are *lists of liquids* that are initially present (rather than being created as a result of following the protocol), and can be regarded as inputs to the protocol. A resource may consist of a single liquid (e.g. if it represents a buffer or reagent), or multiple (e.g. if it represents a set of samples).

The diagram is a node-link representation of a Directed Acyclic Graph representing how particular *lists of liquids* are created from other *lists of liquids* as the protocol is carried out. In this diagram, the nodes represent *lists of liquids*, and the operations that act on them. A *resource* is particular kind of *list of liquid* that may contain a single liquid (e.g. if it represents a buffer), or multiple (e.g. if it represents a set of samples).

The volumes of liquid to be transferred are labeled on the corresponding arrows: this may be a single volume (in which the same volume is taken of all liquids), a list of volumes of the same length as the list of liquids, or, if a node represents a single liquid, a list of volumes of any length.

A pair of *lists of liquids* can be combined in two logical ways: given two lists *x* and *y*, the Cartesian product (cross) independently combines every element of *x* with every element of *y*, whereas the convolution operation (`zip`) combines each element of *x* with only the corresponding element of *y*. If one of the lists consists of only one element, then it is combined with each element of the other list. The operation of combining the lists is represented by a node, labeled as either `zip` or cross. As an example, a `zip` operation can be used to combining pairs of PCR primers, and a `cross` operation can be used to combine each of the resulting mixtures with each of several samples.

The diagram visually distinguishes between different ways in which things can be combined (Figure 2): a solid arrow indicates that the corresponding liquids remained in the same wells, rather than being transferred to new wells. When two liquids are combined, the first could be added to the second (dashed arrow from first, solid arrow from second), the second could be added to the first (dashed arrow from second, solid arrow from first), or both could be added to a new, empty, container (dashed arrows from both).

**Figure 2:**
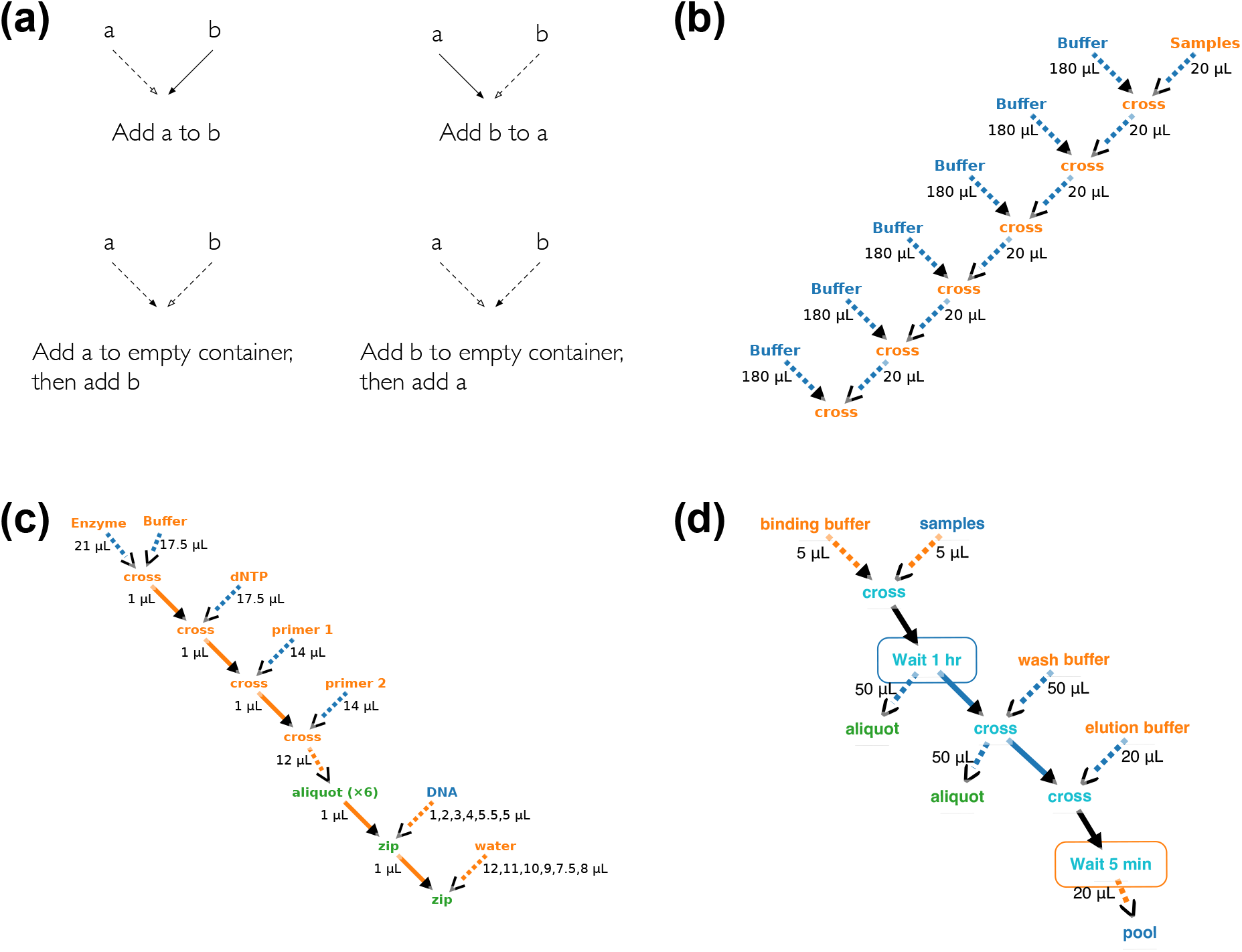
Different arrow styles corresponding to 4 ways of mixing (**a**), and diagrams for 3 example protocols, representing serial dilution (**b**), setting up for a PCR run (**c**), and a SequalPrep extraction (**d**).

## 3 Interface for editing

The graphical interface is implemented as a web-application, providing cross-platform support. The back-end is written in Python using the Flask framework (9); the front-end is written in JavaScript using the D3.js library (10, 11) and cola.js (12) for network layout. Once a user has logged in they are able to create a new protocol or edit a protocol that they have previously created.

### Defining a reference

*Containers, pipettes*, and *resources* can be added by clicking on the ‘add’ link under the appropriate list. Containers and pipettes can be added at any time: before adding resources, as the diagram is being drawn, or after all operations have been added.

### Adding a resource node

When a resource is added, a corresponding node will be added to the diagram. Additional nodes can be created by dragging the name of the resource from the list of resources and dropping it on the diagram.

### Deleting a node

Any node can be deleted by right-clicking on it, and selecting ‘Delete’ from the context-menu. An exception is the final node corresponding to a resource, which cannot be deleted.

### Adding or deleting an operation

The user can click on a node and drag onto another to indicate that they should be combined in a new location; this creates nodes corresponding to the combination operation and the resulting object. If the shift key is held down when the mouse is released, this will be interpreted as meaning that the node that was dragged from should be *added to* the node that was *dragged to*, rather than both being transferred to new wells. Right-clicking on the node representing the operation opens a context menu that allows the combination type to be changed between `zip` and `cross`.

The context menu can also be used to add nodes representing operations that act on only one node, such as performing an operation other than liquid-handling (e.g. thermocycling) or transfering something from one location to another without combining with anything else (taking an aliquot, spreading onto agar, or picking colonies). Some operations act on entire containers, rather than particular wells: this is indicated by a rectangle surrounding the node representing the process. Dragging between two nodes representing operations of the same type acting on lists of liquids located in the same container merges them: they are moved into the same rectangle, and changing the options of one in the panel updates the other. When reading the diagram the nodes should be interpreted as representing the list of liquid resulting from performing the corresponding operation on the inpur list of liquids.

### Editing an operation

Clicking on a node selects it, highlights it, and displays its details in an information panel through which they can be edited. For nodes representing a list of liquids, this displays the contents of each element of the list, which cannot be directly edited. For processing operations, this allows the relevant parameters to be edited (e.g. temperatures and durations for thermo-cycling).

### Editing link details

Clicking on an edge (or its arrowhead, which provides a larger target) selects it and displays its details in the side panel. It is also possible to change the parent of an edge by right-clicking, selecting ‘Change parent’ form the context menu, and then clicking on the node that should become the parent.

### Copying part of a diagram

after clicking on the ‘select nodes to copy’ button, clicking on a node toggles whether it is selected, and multiple nodes can be selected. Clicking on the ‘Copy’ button then copies each of these nodes, and every link that is incident to one of these nodes. This makes it easy to specify that part of a protocol should be repeated, for example with both experimental samples and controls.

### Assigning to containers and wells

The information panel that opens when a node is clicked allows the corresponding container to be selected from a list populated with all the containers that have been defined.

Once a node has been assigned to a container, the specific wells in which it will be located can be set by either clicking on “Show or set well locations” in the information panel for that node, or “Well locations” beside the name of the container in the list of containers. Either of these will open a model window showing the container’s wells on the left, and a list of liquids assigned to the container on the right. Lists of liquids corresponding to a node are grouped together and assigned the same colour; the constituent liquids are displayed as a numbered list, and their components listed.

A single numbered liquid can be dragged from the list and dropped onto a well. Alternatively, a coloured bar can be dragged to simultaneously assign the entire list of liquids corresponding to a node. In the latter case, clicking on the appropriate icon determines whether these will be placed in a row, column, or rectangle.

### Exporting protocols

Above the diagram are a series of links to download the protocol in various formats. If details required to convert the protocol into the desired format are missing (e.g. if a transfer does not have a pipette specified) then a modal dialog will appear listing the errors, and the corresponding parts of the diagram will be highlighted.

The source code, and link to a demo, is available at https://github.com/jamesscottbrown/list-of-liquids.

## Acknowledgement

J.S-B acknowledges funding through the EPSRC & BBSRC Centre for Doctoral Training in Synthetic Biology (EP/L016494/1), and from Dstl. AP acknowledges funding from the EPSRC (EP/M002454/1).

